# The Role of Sevenless in Drosophila R7 Photoreceptor Specification

**DOI:** 10.1101/624155

**Authors:** Andrew Tomlinson, Yannis Emmanuel Mavromatakis, Ronald Arias

## Abstract

Sevenless (Sev) is a Receptor Tyrosine Kinase (RTK) that is required for the specification of the *Drosophila* R7 photoreceptor. Other *Drosophila* photoreceptors are specified by the action of another RTK; the Drosophila EGF Receptor (DER). Why Sev is required specifically in the R7 precursor, and the exact role it plays in the cell’s fate assignment have long remained unclear. Notch (N) signaling plays many roles in R7 specification, one of which is to prevent DER activity from establishing the photoreceptor fate. Our current model of Sev function is that it hyperactivates the RTK pathway in the R7 precursor to overcome in the N-imposed block on photoreceptor specification. From this perspective DER and Sev are viewed as engaging the same transduction machinery, the only difference between them being the level of pathway activation that they induce. To test this model, we generated a Sev/DER chimera in which the intracellular domain of Sev is replaced with that of DER. This chimerical receptor acts indistinguishably from Sev itself; a result that is entirely consistent with the two RTKs sharing identical transduction abilities. A long-standing question in regard to Sev is the function of a hydrophobic domain some 60 amino acids from the initiating Methionine. If this represents a transmembrane domain, it would endow Sev with N-terminal intracellular sequences through which it could engage internal transduction pathways. However, we find that this domain acts as an internal signal peptide, and that there is no Sev N-terminal intracellular domain. *phyllopod* (*phyl*) is the target gene of the RTK pathway, and we show that R7 precursors are selectively lost when *phyl* gene function is mildly compromised, and that other photoreceptors are removed when the gene function is further reduced. This result adds a key piece of evidence for the hyperactivation of the RTK pathway in the R7 precursor. To facilitate the hyperactivation of the RTK pathway, Sev is expressed at high levels. However, when we express DER at the levels at which Sev is expressed, strong gain-of-function effects result, consistent with ligand-independent activation of the receptor. This highlights another key feature of Sev; that it is expressed at high levels yet remains strictly ligand dependent. Finally, we find that activated Sev can rescue R3/4 photoreceptors when their DER function is abrogated. These results are collectively consistent with Sev and DER activating the same transduction machinery, with Sev generating a pathway hyperactivation to overcome the N-imposed block to photoreceptor specification in R7 precursors.

## Introduction

The Drosophila ommatidium is a precisely structured cellular unit containing eight photoreceptors and varied support cells. It has become a classic model for understanding how short-range positional signals direct cells to their fate, and the specification of the R7 photoreceptor has been the focus of many studies. Research initially focused on R7 because of the *sevenless* (*sev*) mutation in which the R7 precursor fails to be specified to its normal fate but is redirected to become a lens-secreting cone cell. Characterization of the gene showed that it encoded a Receptor Tyrosine Kinase (RTK)(Hafen et al., 1987) that was required for the R7 precursor to receive positional signals necessary for the specification of the R7 fate. Thereafter, the role played by another RTK - the Drosophila EGF-Receptor (DER) – was described (Freeman, 1996, Schejter and Shilo, 1989). DER acts in the specification of all the photoreceptors (excepting R8), and Sev and DER engage the Ras/MAPK transduction pathway to promote the removal of Tramtrack (Ttk – a transcription factor that represses the photoreceptor fate)(Simon et al., 1991, Bonfini et al., 1992, Tang et al., 1997, Xiong and Montell, 1993, Li et al., 2002). Since Sev and DER appeared to function in the same way, the question arose as to why Sev was selectively required for R7 specification. Three models were proposed. First, that Sev and DER activated identical transduction pathways but that Sev supplied a much more potent activation, sufficient to specify the R7 rather than the R1-6 photoreceptor fate. Second, that Sev provided another signal in addition to the Ras/MAPK pathway, and that that information was used to direct the R7 rather than the R1-6 fate. Third, that timing of the Ras/MAPK pathway activation was critical for the type of cell specified, and that Sev ensured that the pathway was activate at the required time in the R7 precursor. More recently, the roles played the receptor Notch (N) in R7 specification have been a described (Cooper and Bray, 2000, Tomlinson and Struhl, 2001, Tomlinson et al., 2011), and two of these roles are of importance here. First, N activity opposes the actions of DER signaling to achieve the removal of Ttk. Second, the type of photoreceptor specified is not determined by the actions of the RTK pathway; rather, if a cell removes Ttk and has coincident high N activity it becomes an R7 type, if there is no concurrent N activity then the R1-6 photoreceptor fate results. These two features led to a new model in which the role of Sev is to provide a potent RTK activation solely to overcome the N opposition to Ttk removal. That is, Sev does no more than hyperactivate the DER transduction pathway, and its action has no influence on whether the R7 or R1-6 fates are specified.

The intracellular sequences of an RTK direct which transduction pathway it engages, and if indeed DER and Sev activate identical internal cellular machinery, then a chimerical receptor in which the intracellular sequences of Sev are replaced with those of DER should selectively engage the DER transduction pathway. Indeed, we find that such a chimerical receptor functions in a manner indistinguishable from Sev itself, suggesting that the two RTKs activate the same transduction pathways, and that no information that distinguishes R7 from R1-6 is encoded in Sev transducing activity.

Sev has an unusual N terminus with a hydrophobic domain ∼ 60 residues from its initiating Methionine (Simon et al., 1989). If this region acts as a transmembrane domain (TM), then the N terminus of the protein would be inserted through the plasma membrane and provide N-terminal intracellular sequences through which Sev could engage internal machinery and thereby supply additional information to the R7 precursor. However, we find that this domain acts as an internally-displaced signal peptide, and that sequences N-terminal to it play no significant role in Sev function. The chimerical receptor result suggests that DER and Sev engage the same transduction machinery but does not inform on the other aspect of the model; that the RTK pathway is hyperactivated in the R7 precursor. A number of results suggest that the pathway is indeed a hyperactive in R7 (see Discussion), but no evidence was available with regard to *phyllopod* (*phyl*); the target gene of the RTK pathway that encodes an adaptor protein that promotes the removal of Ttk (Tang et al., 1997, Dickson et al., 1995, Li et al., 2002). We show that when *phyl* function is mildly abrogated, only R7s are affected. But when that abrogation is increased then R1-6 class photoreceptors specifications are disturbed.

To facilitate its hyperactivation of the RTK transduction pathway, Sev is expressed at high levels. This contrasts with DER which is expressed at low levels. Furthermore, when we expressed DER at the levels at which Sev is expressed, we observed ectopic activation of the RTK pathway as evidenced by the transformation of cone cell precursors into R7s. This highlights another key feature of Sev; it is expressed at high levels and yet remains strictly ligand dependent. The results from the chimerical protein suggest that this resistance to ligand-independent activation maps to the Sev extracellular domain.

Finally, we show that an activated form of Sev (Basler et al., 1991) can functionally substitute for DER in the specification of R3/4 photoreceptors. This result further supports the uniformity of function of Sev and DER.

## Materials and Methods

### Immunohistochemistry and histology

Protocols for adult eye sectioning and eye disc immunostaining as previously described (Tomlinson and Ready, 1987, Tomlinson et al., 2011). Primary antibodies: Sev.ICD, Myc (Santa Cruz Biotechnology); DER.ICD (Sigma E3031); GFP (molecular Probes); Elav (DSHB); Svp (gift Y.Hiromi); Runt (gift J.Reinitz).

### fly stocks

*sevd2, sev.Gal4* as previously described (Tomlinson et al., 2011, Mavromatakis and Tomlinson, 2013). *UAS.phylRNAi* [GD12579] VDRC. *UAS.derDN* BDSC 5364.

### Generation of transgenes

All transgenes under *sev* transcriptional control were generated and transformed as previously described (Mavromatakis and Tomlinson, 2013). All were cloned in the same *attB* vector and transformed into the same *attP* site (86F) to ensure uniformity of expression. An AvrII site we created immediately 5’ to the sequence encoding the *sev* TM domain, and a NotI site was created immediately 3’. These sites were used to facilitate the cloning of the chimerical receptors. PCR was used to polymerize the appropriate pieces of DNA for cloning into the relevant sites. For the N-terminal protein modifications, an SpeI site was created immediately 5’ to the N-terminal hydrophobic domain to facilitate the insertion of GFP. To substitute the hydrophobic domain with a heterologous signal peptide, a BglII site was created immediately 3’ to the hydrophobic domain, and as previously described (Basler et al., 1991) the Drosophila cuticle protein CP3 signal sequence followed the sequences encoding c-Myc were substituted for sequence lying 5’ to the hydrophobic domain. UAS.sev* was constructed by PCR of the sev* coding sequence then cloned into a *UAS attB* vector and transformed into attP 86F.

## Results

To evaluate whether Sev and DER engage the same transduction pathways, we assayed the signaling abilities of chimeras constructed from the two receptors. Before doing this we performed control validation experiments.

### Validation Experiments

Following the work of Basler et al. (1991), we generated and transformed a construct in which the *sev* coding sequence is expressed under the control of the *sev* enhancer and promoter sequences. When a single copy of this transgene was introduced into a *sev°* background (Fig.1C), 100% of ommatidia (N = 342) had R7s (as opposed to the *sev°* background in which there are zero R7s – Fig.1B). Furthermore, staining with the Sev antibody (raised against the Sev C-terminus) showed a wild type expression pattern in developing third instar discs (Fig.1E), and staining of these discs with Elav (which highlights photoreceptors), Seven-Up (Svp; expressed in R3/4s and R1/6s) and Runt (expressed in R8 and R7s) showed that the R7s were specified at the correct time and place in a manner indistinguishable from wild type eye discs (Fig.1F). This confirms the efficacy of the approach for evaluating the *sev°*-rescuing abilities of chimerical receptors. Next, we generated and transformed a similar construct in which the *der* coding sequence was expressed under *sev* transcriptional control (*sev.DER*). In *sev.DER* eye discs, strong DER antibody (raised against the DER C-terminus) staining was detected in the Sev-expressing cells (Fig.1H) superimposed on the normal low DER expression pattern (Fig.2D). This high-level DER expression caused multiple R7s in adult ommatidia (red arrows in Fig.1G) resulting from the inappropriate specification of cone cell precursors as R7s (Fig.1I). The re-specification of cone cell precursors as R7s occurs when their RTK pathway is hyperactivated (Basler et al., 1991). This highlights the propensity of DER to autoactivate when expressed at high levels (Schweitzer et al., 1995). Furthermore, it contrasts strikingly with Sev that is expressed at high levels to facilitate its potent activation of the RTK pathway, and yet remains strictly ligand dependent.

**Figure 1:**
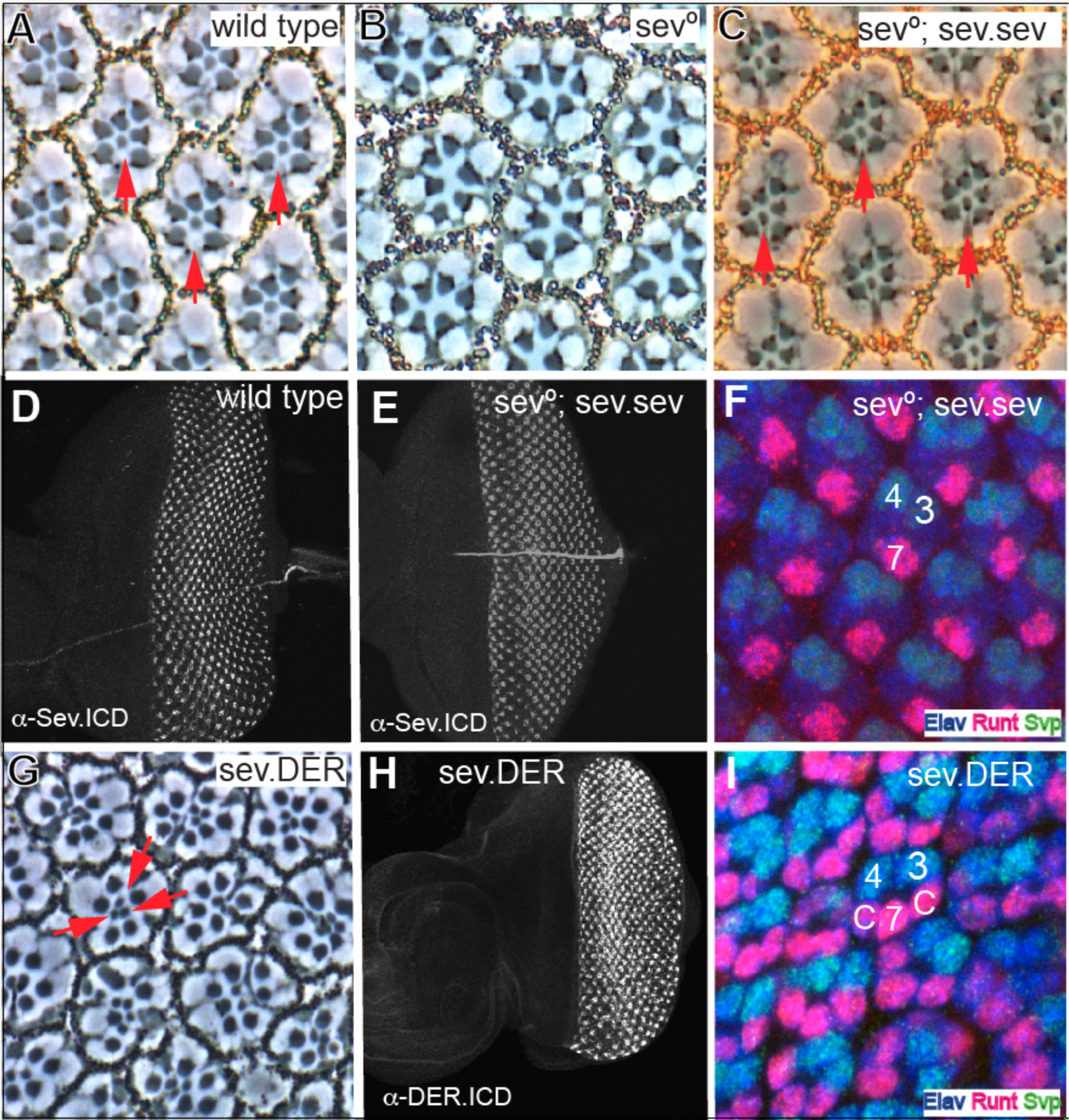
Rescue of *sev°* by Sev expression, and the effects of DER expressed at Sev levels. (**A**) Phase contrast image through a wild type retina in which the rhabdomeres of six outer photoreceptors are arrayed in an asymmetric trapezoid shape. In the center of the array lies the small central rhabdomere of R7 (red arrows). **(B)** R7 is absent from every ommatidium in *sev°* eyes. **(C)** *sev* expressed under *sev* transcriptional control (*sev.sev*) restores R7s (red arrows) to all *sev°* ommatidia. **(D)** An antibody raised against the C-terminus of Sev highlights the typical expression pattern in wild type eye discs. **(E)** *α*-Sev.ICD staining of *sev°; sev.sev* eye discs show the normal Sev expression pattern. **(F)** In *sev°; sev.sev* eye discs, normal pattern formation is evident with R7 specified at the correct time and place. At the two-cone-cell-stage, R7s (highlighted by Elav (blue) and Runt (red) expression) are seen on the other side of the ommatidium from R3/4 labeled by Elav and Svp (green). **(G)** In *sev.DER* adult eyes, many R7-like cells (red arrows) are evident in each ommatidium. **(H)** A *sev.DER* eye disc labeled with an antibody raised against the C-terminal region of DER. Strong DER staining is observed in the Sev-expression cells superimposed upon the normal DER expression pattern (see Fig.2D). **(I)** At the two-cone-cell-stage, sev.DER eye discs show cells in cone cell positions (c) differentiating as R7-like photoreceptors (expressing Runt and Elav).

### The DER ICD substitutes for that of Sev

We next generated and transformed a construct in which the ICD of Sev was replaced with the equivalent domain of DER, and expressed under *sev* transcriptional control (this has the Sev ECD and TM domains and the DER ICD and we refer to it as *sev.SSD* (Figure 3F)). This was introduced into a *sev°* background. Staining with the DER-ICD antibody detected high levels of DER protein in the Sev-expressing cells (Fig.2E,F) superimposed on the normal DER expression (Fig.2D). A single copy of this transgene fully substituted for Sev as evidenced by the complete rescue of R7s in a *sev°* background (Fig.2A, N = 263), and by the specification of R7s at the normal time and place in the developing ommatidia (Fig.2C). When the transgene number was doubled (in *wildtype* or *sev°* background) the phenotypes remained the same, with no evidence of any gain-of-function phenotypes. These results argue that a chimera containing all the intracellular sequences of DER acts as a receptor that is indistinguishable from Sev itself.

**Figure 2:**
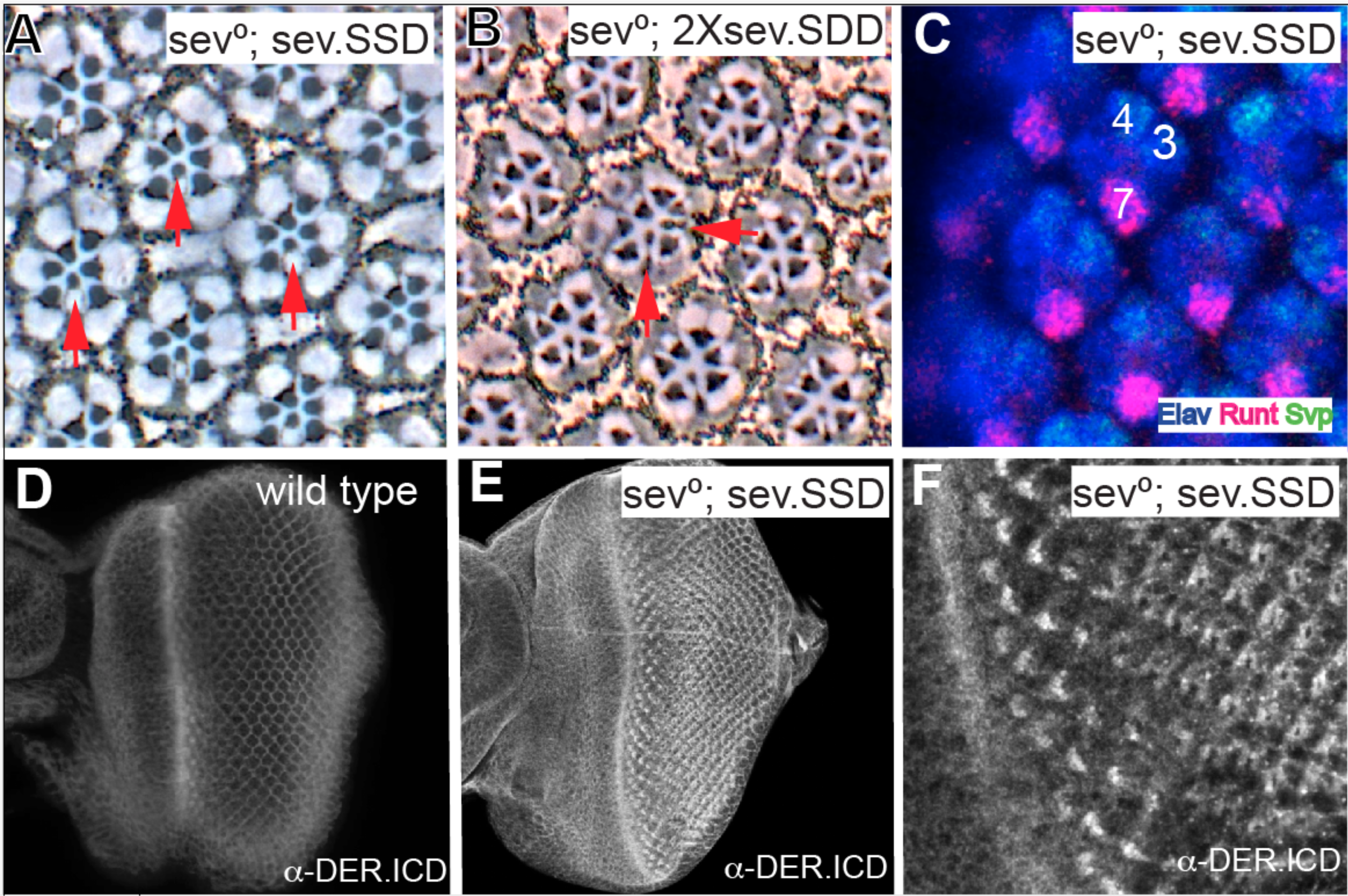
The signaling abilities of Sev/DER chimerical receptors. The *sev.SSD* chimera (Sev ICD replaced with that of DER and expressed under sev transcriptional control) rescues R7s in all *sev°* ommatidia in adult eyes. (B) When the DER TM domain was additionally substituted and expressed under *sev* transcriptional control (*sev.SDD*) in two copies in *sev°*, occasional supernumerary R7s are detected (horizontal red arrow). (C) *sev.SSD* rescues R7s specification at the correct time and place in *sev°* eye discs. (D) A wild type eye disc stained with anti-DER.ICD shows staining in all cells with an upregulation in cells in and behind the morphogentic furrow. (E) *sev°; sev.SSD* eye discs stained with *α* -DER.ICD show strong upregulation in the Sev-expressing cells. (F) High magnification from image in E showing DER expression in the normal Sev pattern.

**Figure 3:**
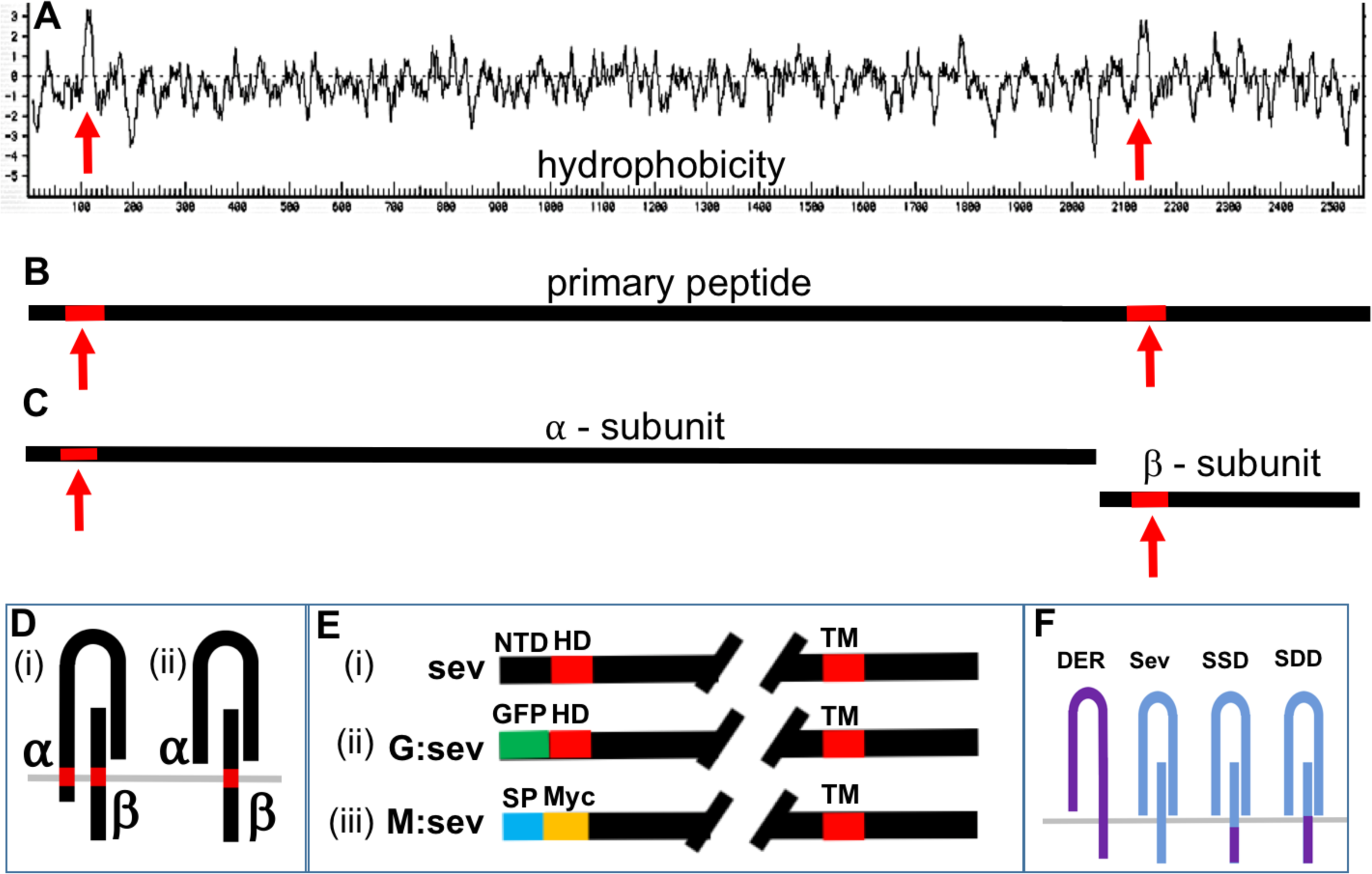
Schematic summary of the molecular features of Sev. (A) A Kyte-Doolittle hydrophobicity plot of Sev reveals two salient hydrophobic domains (red arrows). The one to the right indicates the conventional TM domain. The one to the left may be an N-terminal TM or an internal signal peptide. (B) Shows the Sev primary peptide carrying the two highlighted hydrophobic domains. (C) The primary peptide is cleaved into *α* and *β* subunits. (D) Two topological arrangements of the *α* /*β* dimer: (i) if the N-terminal hydrophobic domain is a TM, there is an N-terminal ICD; (ii) if it represents and internal signal peptide, all *α* subunit sequences are exclusively extracellular. (E) Visual description of the N-terminal architecture of three transgenes. In *sev.sev*, ∼60 residues (black) lie N-terminal to the hydrophobic domain (HD – red). In *sev.G.sev* the N-terminal ∼60 amino acids are replaced by GFP (green). In *sev.M.sev* the N-terminal residues are removed, and the HD domain is replaced with a conventional signal sequence (blue) followed by a Myc tag (yellow). (F) Schematic description of DER and Sev and the two chimeras (*SSD* and *SDD*).

Next, we generated and transformed a construct in which the TM and ICD of Sev were replaced by those of DER (*sev.SDD*; Fig.3F). When introduced into a *sev°* mutant background, one copy of this transgene rescued R7s in all ommatidia (not shown, N = 246), but when the transgene number was doubled, supernumerary R7s (Fig.2B; red arrows) were evident at a low frequency (∼1% of ommatidia), suggesting that the protein has a slight tendency to autoactivate when expressed at high level.

### The role of the Sev N-terminal region

Work from Simon et al., (1989) highlighted two features of Sev. First, that the primary peptide is cleaved into *α* and *β* subunits that form dimeric structures (Fig.3B-D) which are then likely assembled into a tetramer, similar in overall organization to the Insulin Receptor. Second, and particularly germane to this work is the presence of an N-terminal hydrophobic domain some 60 residues from the start of the protein (Fig.3A). If this domain represents a TM domain, then the *α–*subunit will be a Type II transmembrane protein with an N-terminal ICD (Fig.3D(i)). If this is the case, then *sev.SSD* transgene will not replace all Sev intracellular sequences. If, however, the N-terminal hydrophobic domain represents an internal signal peptide, then the N terminal 60 residues would be cleaved and lost from the receptor (Fig.3D(ii)). To resolve these Sev topological issues we generated and transformed two additional transgenes.

#### (i) Replacement of the Sev N terminal region with GFP

We replaced the residues N-terminal to the hydrophobic domain with GFP (*sev.G:sev*; Fig.3E(ii))). A single copy of this transgene in a *sev°* background rescued R7s in ∼65% of ommatidia (Fig.4A). In the presence of two copies of this transgene, rescue of R7s was 100% (N = 174). *α*-GFP staining of *sev.G:sev* in *sev°* eye discs showed GFP expression in the Sev-expressing cells (Fig.4D), but only weak Sev staining was evident using *α*–Sev.ICD (Fig.4E). To evaluate this low level of Sev expression, we co-incubated *sev.sev* eye discs in the same tubes. Fig.4F shows the higher level of Sev staining expressed by these control discs. Thus, when the N terminal region of Sev is replaced with GFP, there is a reduction in the level of the expression of the receptor that correlates with its reduced ability to rescue *sev°*. But, when the transgene number was doubled, then a complete rescue occurred (Fig. 4B,C), which argues against the ∼ 60 N-terminal residues endowing Sev with any critically-required signaling abilities. Rather we infer that the GFP substitution results in decreased amounts of functional receptor. When we increased the confocal laser power, we observed the *α*-GFP and *α*–Sev.ICD in different parts of the cells, indicating that they did not necessarily co-localize to a single protein. This is consistent with the N-terminal hydrophobic domain acting as a signal sequence resulting in GFP being cleaved from the protein.

**Figure 4:**
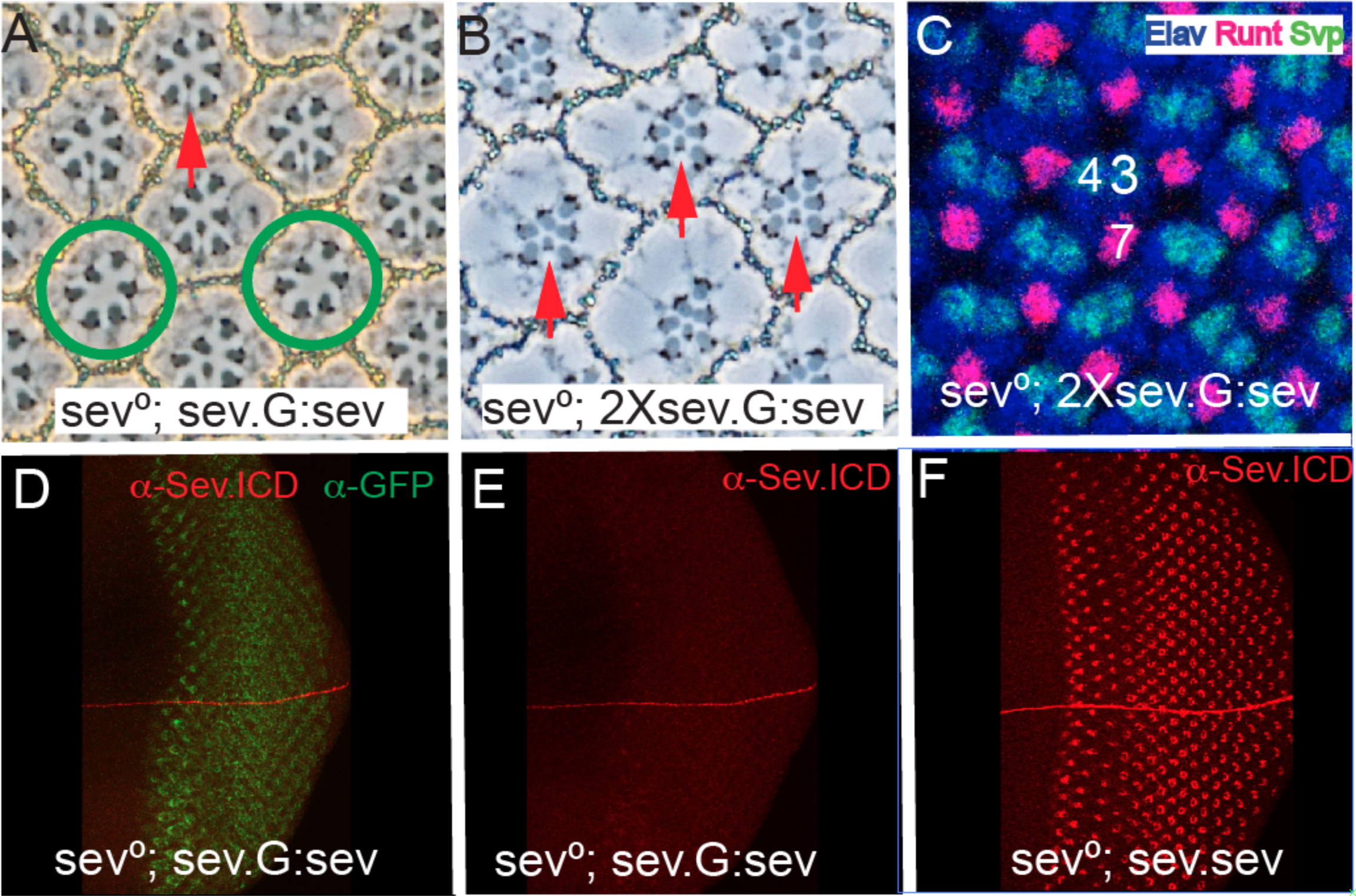
The effects of substituting the Sev N-terminal ∼60 residues with GFP. (A) When the Sev N-terminal ∼60 amino acids are replaced with GFP and expressed under sev transcriptional control (*sev-G:sev*) one copy of the transgene fails to rescue all R7s of *sev°* ommatidia. Red arrow indicates a rescued R7, and the green circles highlight ommatidia lacking an R7. (B) Two copies of *sev-G:sev* rescue R7s in all *sev°* ommatidia. (C) Two copies of *sev-G:sev* rescue R7s at the correct time and place in *sev°* eye discs as evidenced by Runt and Elav positive nuclei in the cells occupying the R7 positions. (D) GFP staining of sev°; *sev-G:sev* eye discs shows expression in the Sev-expressing cells. (E) *α*-Sev.ICD staining of sev°; *sev-G:sev* shows only weak staining. (F) The staining level of *α*-Sev.ICD in sev° sev.sev eye discs co-incubated with sev°; *sev-G:sev* eye discs shown in D.

#### (ii) Replacement of the N terminal hydrophobic region with a heterologous signal peptide

To investigate further whether the N terminal hydrophobic region acts as an internal signal peptide, we removed the N terminal region and replaced the hydrophobic stretch with a conventional signal peptide followed by a Myc tag to label the N terminus of the protein (Fig.3E(iii)). This construct (*sev.M:sev*) was transformed, and in a *sev°* background a single copy rescued R7s in all adult ommatidia (Fig.5A) (N = 187). Pattern formation occurred normally in these developing eye discs (Fig.5B,C) in which R7s were specified in the correct time and place. When *sev°; sev.M:sev* discs were stained with *α*-Myc and *α*-Sev.ICD, both showed the normal Sev-expression pattern, and both were coincidently localized (Fig.5D-F). Thus, the removal of the N-terminal ∼60 amino acids, and the replacement of the N-terminal hydrophobic domain with a heterologous signal peptide results in a protein that fully substitutes for Sev. This result suggests that the N-terminal hydrophobic domain in Sev acts as signal peptide and that there are no N-terminal intracellular sequences in Sev.

**Figure 5:**
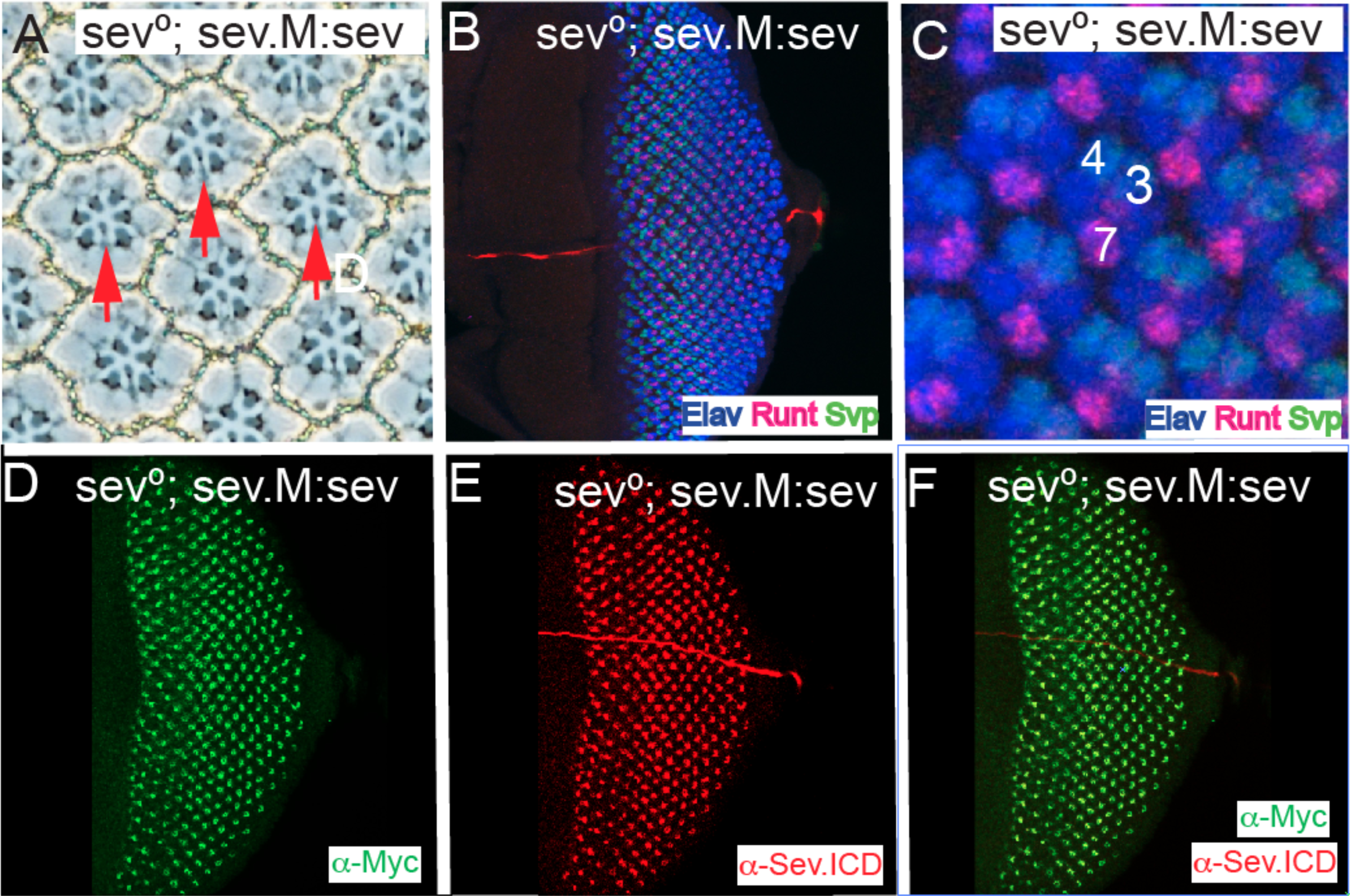
A heterologous signal sequence substitutes for the Sev N-terminal hydrophobic domain. All panels show images from *sev°; sev.M:sev* animals in which the N-terminal hydrophobic domain and all residues N-terminal to it are replaced with a conventional signal peptide followed by the Myc tag. (A) In adult eyes all R7s are rescued (red arrows). (B-F) Eye discs stainings. (B, C) show that pattern formation is normal as monitored by Elav, Runt and Svp expressions, and R7s are specified at the correct time and position. (D) *α*-Myc staining highlights the normal Sev expression pattern. (E) *α*-Sev.ICD shows the normal Sev expression pattern. (F) Superimposition of the Myc and Sev staining show their coincident expression in the Sev expressing cells.

### Combining the N and C-terminal modifications in a single transgene

The results to date suggested that the ICD of Sev could be functionally substituted by the equivalent domain of DER, and that the N-terminal domain of Sev was unlikely to be an ICD. To control for the unlikely possibility that redundancy existed between the N and C termini of Sev in determining which intracellular pathways are activated, we next generated a chimera containing both the N- and C-terminal modifications. Specifically, we generated and transformed a construct in which the heterologous signal peptide is present at the N-terminus (and endowing the Myc tag), and the ICD of Sev is replaced with that of DER (*sev.M:SSD*). When introduced into a *sev°* background, a single copy of this transgene rescued the R7s of all ommatidia (Fig.6A; N = 262); pattern formation in developing third instar disc appeared fully wild type (Fig.6B) with R7s specified at the correct time and place (Fig.6C). When these discs were stained with *α*-Myc, the normal Sev-expression pattern was detected (Fig.6D), and the DER-ICD antibody highlighted the expression in the Sev-expression cells superimposed on the normal DER expression pattern (Fig.6E). In these eye discs the Myc and DER antibodies were found colocalized in the Sev-expressing cells of the eye disc (Fig.6F) which is consistent with them recognizing the same protein.

**Figure 6:**
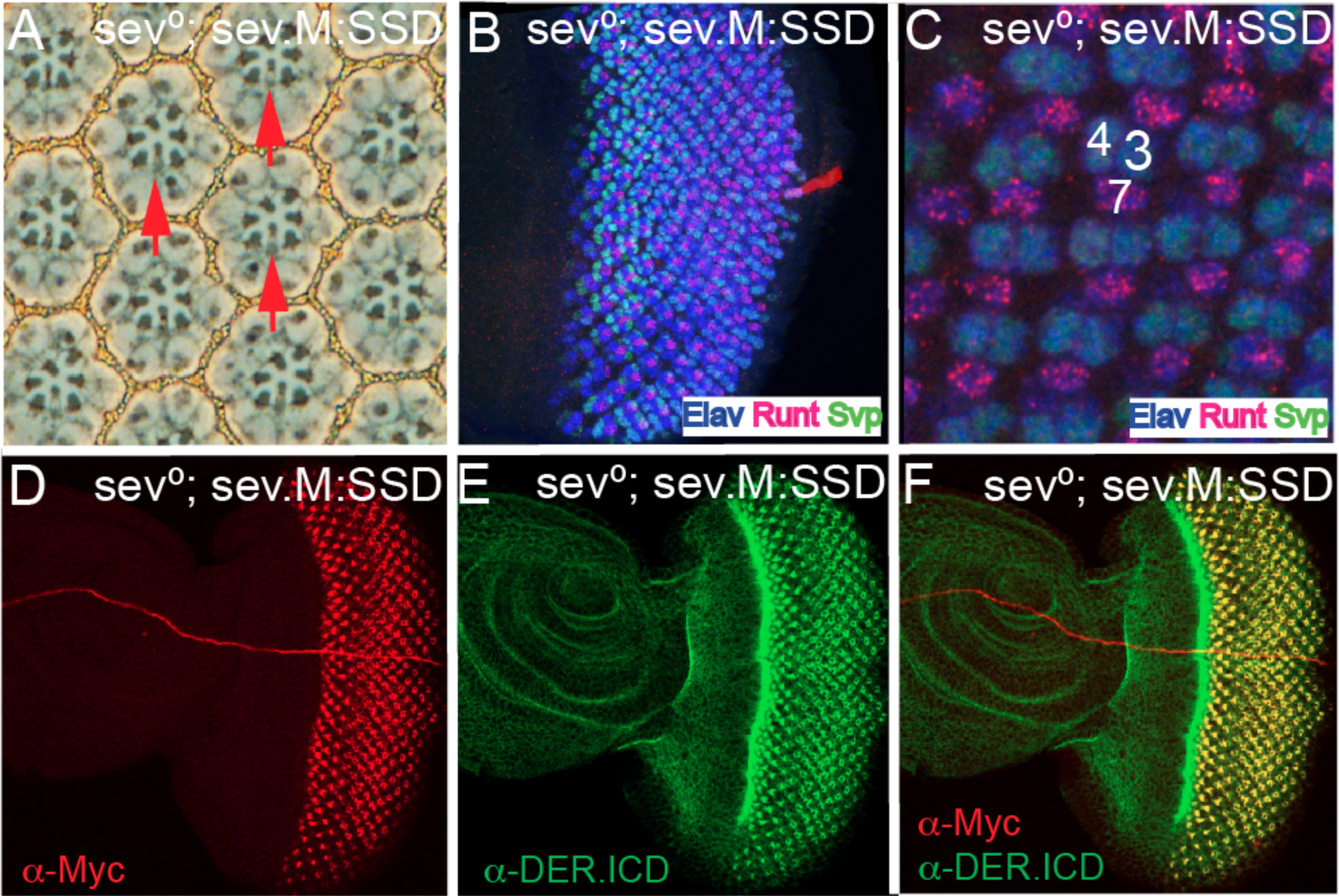
A chimera with the DER ICD and a heterologous signal sequence rescues sev°. All panels are from *sev°; sev.M:sev* animals in which the N-terminal hydrophobic domain and all residues N-terminal to it are replaced with a conventional signal peptide followed by the Myc tag, and the Sev ICD is replaced with that of DER. (A) In adult eyes all R7s are rescued (red arrows). (B-F) Eye discs stainings. (B, C) show that pattern formation is normal as monitored by Elav, Runt and Svp expressions, and R7s are specified at the correct time and position. (D) *α*–Myc staining highlights the normal Sev expression pattern. (E) *α*–DER.ICD shows the Sev expression pattern superimposed on the normal DER expression. (F) Superimposition of the Myc and DER staining show their coincident expression in the Sev expressing cells.

### Abrogation of *phyl* gene function preferentially affects R7s

*phyl* is the RTK target gene in R1/6 and R7 precursors, and if indeed the pathway is hyperactive in R7s then they may show differential sensitivity to reductions in *phyl* gene function. To investigate this, we drove expression of *UAS.phylRNAi* using *GMR.Gal4* (which is expressed ubiquitously in all the developing ommatidia). When a single *RNAi* transgene was present, R7s were preferentially lost from adult ommatidia (Fig.7A shows the typical *sev°* phenotype in which only R1-6 cells are present), but when two copies were present, ommatidia containing only four large rhabdomere photoreceptors were present (Fig.7B) which likely results from the additional loss of the R1/6 cells. Thus, reduction of *phyl* gene function preferentially affects R7 precursors; a result consistent with need for a stronger RTK signal transduction in these cells.

**Figure 7:**
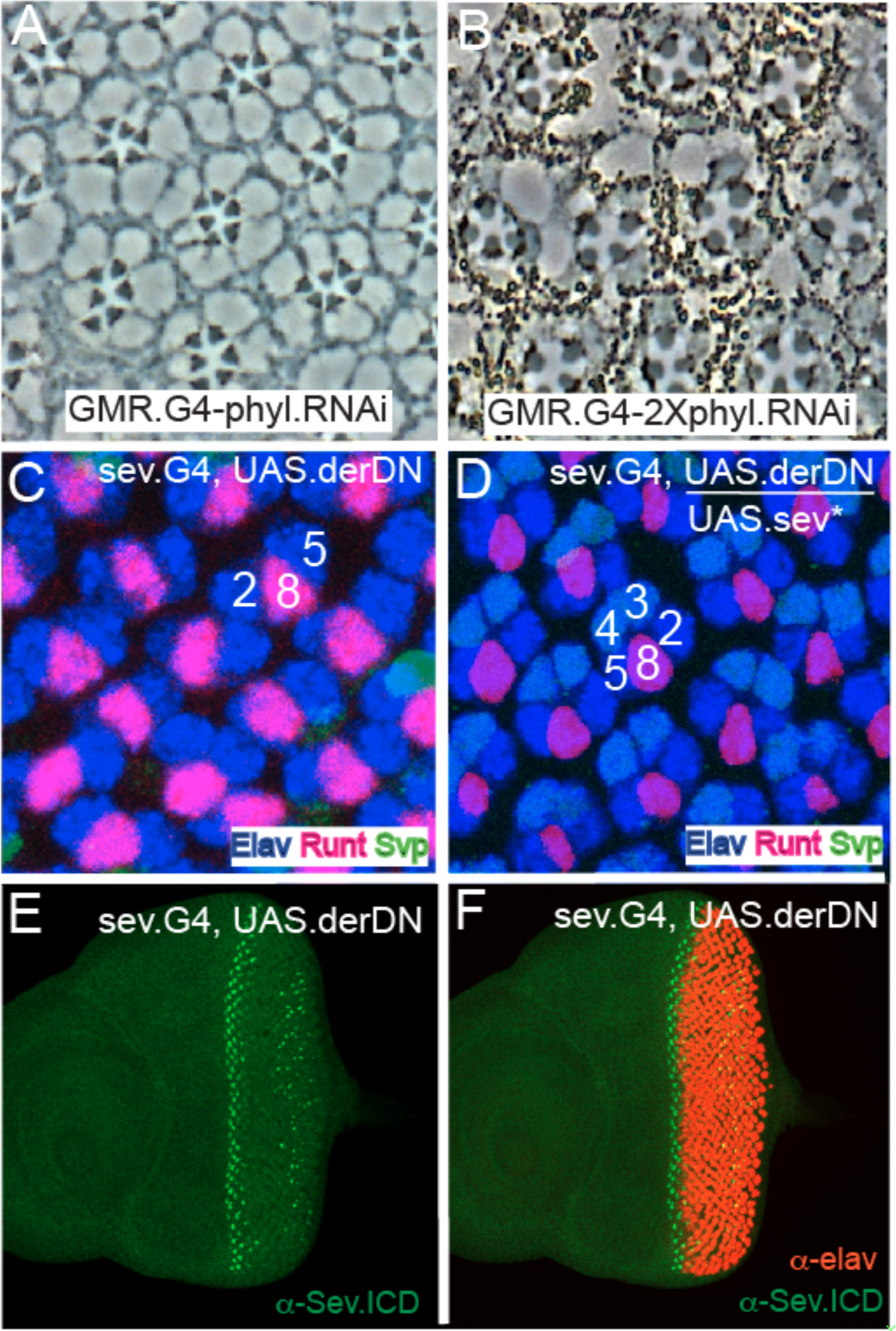
Selective R7 sensitivity to *phyl* gene function reduction, and Sev* rescue of R3/4 DER function. (A) Section through a *GMR.Gal4; UAS.phyl* adult eye shows the typical *sev°* phenotype with each ommatidium specifically lacking R7. (B) When *phyl* gene function is further compromised (*GMR.Gal4; 2XUAS.phyl.RNAi*) only 4 large rhabdomere photoreceptors are evident in most ommatidia. (C) *sev.Gal4; UAS.derDN* eye discs stained for Elav (blue) Svp (green) and Runt (red). Only R2/8/5 clusters are evident indicating that photoreceptor recruitment ceases at this stage. (D) When *UAS.sev*\* is introduced into the *sev.Gal4; UAS.derDN* background, the recruitment of the R3/4 photoreceptors is rescued. (E) *α –*Sev staining (green) indicates normal expression in R3/4 precursors in the early ommatidia, but is degenerate in later ommatidia. (F) Co-staining of *sev.Gal4; UAS.derDN* with Sev and Elav highlights the normal R3/4 Sev expression before photoreceptor differentiation begins.

### Rescue of DER function by Sev

The collective results argue that the ICD of Sev can be functionally substituted by the equivalent domain of DER, and suggests that Sev engages the same transduction machinery as DER. This begs the question of whether the ICD of DER can be functionally replaced by the equivalent domain of Sev. Preliminary experiments to determine whether a transgene ubiquitously expressing DER can rescue *der°* clones failed; we were unable to find a level of DER expression that would rescue the loss of function but not be too high to trigger autoactivation. This prevented the evaluation of which DER/Sev chimeras could rescue DER function. We therefore adopted a different approach. First, we abrogated DER function in R3/4 precursors by driving the expression of a dominant negative form of DER (*UAS.derDN*) with *sev.Gal4* (*sev-derDN*). *α*-Sev-ICD staining showed the normal expression pattern in the R3/4 cells of *sev-derDN* eye discs, but an aberrant pattern in more mature ommatidia where sporadic Sev-positive cells were detected in the posterior portions of the disc (Fig.7E,F). When we examined ommatidial patterning in these discs using standard cell fate markers, we found that the ommatidial progression largely “froze” at the R2/8/5 stage; R3/4 cells were only rarely recruited to the cluster (Fig.7E,F). Second, we introduced a *UAS.sev*\* (a transgene in which activated *sev* is expressed under *UAS* transcriptional control) into the *sev-derDN* background and observed developing ommatidia in which differentiating R3/4 cells were restored (Fig.7D). sev* encodes a form of Sev in which the majority of the ECD is lost but the TM and ICD domains are normal (Basler et al., 1991). Since the expression of this protein rescues *sev-derDN* R3/4s this suggests that the Sev ICD can functionally substitute for DER in these cells. *sev.derDN* eyes have severely deranged adult eyes, which prevented assessment of the loss and rescue of R3/4s in adult ommatidia.

## Discussion

We document here that the SSD chimerical receptor fully rescues *sev°*. That is, this chimera acts in a manner indistinguishable from Sev itself. Since the ICD of an RTK normally directs which transduction pathway a receptor engages, this result argues that in the context of R1/6 and R7 specifications, Sev and DER activate the same intracellular machinery. Originally, there were three different models of the role played by Sev in R7 specification, and each of them is reviewed below in the light of this new result.

### Models that postulated hyperactivation of the RTK pathway

Original hyperactivation models envisioned that the potent transduction of the RTK pathway in the R7 precursor would promote the expression of genes that the lower levels of transduction in R1/6 precursors would not, and that those gene expressions would play key roles in specifying the R7 versus R1/6 fate. The current model argues that the hyperactivation of the RTK pathway in the R7 precursor acts solely to counteract the N-imposed block and ensure that Ttk degradation occur. The rescue of *sev°* by *sev.SSD* is consistent with both these models.

#### Evidence for RTK pathway hyperactivation

A number of pieces of evidence suggest that the RTK pathway is hyperactivated in the R7 precursor: (i) Rap is required in concert with Ras to transduce the RTK pathway in R7s but in R1/6 precursors Ras alone supports the pathway transduction (Mavromatakis and Tomlinson, 2012). This suggests that the Ras alone is insufficient to mediate the pathway hyperactivation in R7 precursors and Rap is additionally engaged to mediate the extra “traffic”. (ii) Levels of MAPK phosphorylation (staining with *α*-dpERK) are high in R7 and not detected in R1/6 precursors, suggesting that MAPK activation occurs at a much higher level in R7s (Kumar et al., 1998). (iii) Sina is the Ubiquitin Ligase that Ubiquitinates and targets Ttk for degradation. In *sina°* eyes, R7s are selectively lost while there are only minor effects on R1/6 (Carthew and Rubin, 1990), suggesting that Sina plays only a minor role in R1/6 photoreceptor specification, but is critically required for Ttk removal in R7 specification.

In this work we examine the sensitivities of the different photoreceptor precursors to the level of *phyl* gene function. *phyl* is the target gene of the RTK pathway in both types of photoreceptor precursors, and yet the R7s are clearly more sensitive to the abrogation of *phyl* gene function than are its R1/6 counterparts. This observation suggests that R1/6 precursors require a lower expression of *phyl* than the presumptive R7s and is consistent with the view that the RTK transduction pathway is hyperactive in the R7 precursor.

#### The role of N in distinguishing R7 from R1/6

It is important to note that in all the manipulations described above that highlight the hyperactivation of the RTK pathway in R7 precursors, none result in a switch from the R7 to the R1/6 fate (or vice versa), all result in the failure to remove Ttk and the resulting specification of the non-neural cone cell fate. Thus, there is no evidence that the hyperactivation of the RTK pathway mediates any information that distinguishes the R7 from the R1/6 fate. Rather, we argue that the type of photoreceptor specified is controlled exclusively by N signaling. That is, when Ttk is removed (the generic photoreceptor fate) and N activity is high, then the R7 type photoreceptor is specified, whereas when N function is low, then the R1/6 class photoreceptor results

### Evidence against Sev activating additional transduction pathways

Earlier models of Sev function envisioned that it activates other transduction pathways in addition to the MAPK cascade to supply extra information to the R7 precursor up-and-above that supplied by DER. In this work we have exchanged all Sev intracellular sequences with topologically equivalent sequence from DER, and find that the resulting chimera functions in a manner indistinguishable from Sev itself. This does not formally exclude the possibility that Sev engages additional transduction pathways; transmembrane proteins that associate with extracellular portions of Sev could mediate the activation of the additional pathways. However, in the context of how we understand RTKs to function, and given all the other information pertaining to Sev, Occamist reasoning argues that Sev and DER activate the same cellular transduction machinery.

### Evidence against the temporal model of cell fate specification

The temporal model envisioned that it was the timing of the MAPK pathway activation that provided the critical information that determined the type of cell specified. This emerged before an appreciation of the roles played by N was achieved, at a time when the RTK pathway alone appeared to control all the sub-type identities of the ommatidial cells. The evidence against this model is now strong. Consider the R1/6, R7 and cone cell specifications which occur in a defined time progression. If the RTK/N signaling information is altered in these cells, their fates can be cleanly inter-switched without affecting the timing of the events (Tomlinson et al., 2011).

### The structure of Sev

The N-terminal hydrophobic domain of Sev has been enigmatic for some time; it was not known whether it acted as an internal signal peptide or a TM domain. If it indeed acted as a Type II membrane anchoring domain, then the ∼ 60 residues that lie N-terminal to it would be intracellular sequences through which Sev might engage internal machinery. Our results argue otherwise, indicating that the N-terminal hydrophobic domain is a signal peptide.The domain can be replaced with heterologous signal peptide (in which the ∼60 residues N-terminal to it are deleted) and a fully-functional and appropriately expressed Sev protein is produced. A similar internal signal peptide is found in the Drosophila Insulin Receptor (Fernandez et al., 1995), and the reason why the domain is displaced from the N terminus of the protein is unknown. Interestingly, this domain is conserved in all 12 members of the sequenced Drosophila phylogeny(Drosophila 12 Genomes et al., 2007), but Diptera outside this group do not have it, and the role it plays remains enigmatic.

### The resistance of Sev to ligand-independent activation

The DER class RTKs are monomeric structures that dimerize (and thereby activate) in the presence of ligand. When such receptors are expressed at high levels, dimerization occurs spontaneously leading to ligand-independent activity. This feature is evident when DER is expressed at high levels under *sev* transcriptional control. Here, the ectopic activation of the RTK pathway liberally re-specifies many cells as photoreceptors (Fig.1G). This contrasts strikingly with Sev, which when expressed at those same levels remains strictly ligand-dependent. The analysis of the chimeras suggests that the resistance of Sev to autoactivation maps to its ECD. When both the TM and ICD of Sev were replaced with the equivalent domains of DER, a low frequency of supernumerary photoreceptors resulted. This occurred in ∼1% of ommatidia and represents only a minor effect. The cleavage of the ECD into *α* and *β* subunits, and their assembly into an InR-like tetrameric structure would account for the resistance of Sev to autoactivation. Presentation of ligand to this class of RTK does not promote oligomerization and activation (the sub-units are preassembled into the tetramers) rather, ligand binding triggers allosteric changes in the receptor that promotes activation.

